# CasLocusAnno: a web-based server for annotating Cas loci and their corresponding (sub)types

**DOI:** 10.1101/459131

**Authors:** Chuan Dong, Zhi Zeng, Qing-Feng Wen, Shuo Liu, Meng-Ze Du, Yi-Zhou Gao, Zhen Liao, Nicholas Saber, Jian Huang, Feng-Biao Guo

## Abstract

CRISPR-Cas systems are prevalent in bacterial and archaeal genomes, and these systems provide a powerful adaptive immune system against predation by phages and other mobile genetic elements (MGEs). They also contribute to other functions, such as gene regulation in prokaryotic organisms. Determining Cas proteins and Cas loci can help mine Cas proteins and facilitate the identification of Cas-associated accessory proteins. Therefore, the purpose of this work is to develop a web-based server, CasLocusAnno, to annotate Cas proteins and Cas loci and to classify them according to (sub)type based on a previous study. CasLocusAnno can annotate Cas proteins and Cas loci and assign their (sub)types within ∽28 seconds for whole protein sequence submissions, with protein sequence numbers ranging from ∽30 to ∽10500. Comparison with Makarova *et al.*’s benchmark data demonstrates that CasLocusAnno can accurately identify Cas loci and (sub)types. In addition, CasLocusAnno can identify Cas proteins with higher accuracy and a lower additional prediction rate (APR) than two excellent software programs, CRISPRCasFinder and MacSyFinder. The domain alignment of a Cas protein can be easily browsed in the annotation results. Our server can be freely accessed at http://cefg.uestc.edu.cn/CasLocusAnno/.

## INTRODUCTION

CRISPR-Cas is short for “<u>c</u>lustered, <u>r</u>egularly <u>i</u>nterspaced, short, palindromic repeats (CRISPR) and <u>C</u>RISPR-<u>as</u>sociated proteins (Cas)”. CRISPR-Cas systems can be categorized into two classes according to the number of subunits used by the effector protein: class 1, encompassing types I, III, and IV, and class 2, encompassing types II, V, and VI (1,2). The former employs multisubunit Cas effectors for immune activity, whereas the latter adopts a single effector protein with different functional domains, such as Cas9, which possesses HNH and Ruvc domains, and Cpf1 (Cas12a), which possesses only an Ruvc-like domain, to engage in interference (3). These six types can be further divided into several subtypes according to their signature proteins and the architectural features of the Cas locus (1,2). The diverse CRISPR-Cas systems mainly provide powerful adaptive immune activity for Bacteria and Archaea against predation by invaders such as phages, plasmids and other parasites. The general outline of how CRISPR-Cas systems carry out their immune functions has been presented by research communities. Their defense activity consists of three steps (4–6): first, some DNA segments derived from MGEs are inserted into CRISPR arrays to become spacers with the help of part of Cas proteins in the adaptation stage, and the spacer sequence can function as markers, which record memory of the invaders; second, as a functional element, the CRISPR array is transcribed into precursor CRISPR-associated RNAs, which become crRNAs and are further armed with Cas effector protein(s) in the crRNA maturation and Cas protein assembly stage; third, crRNA-Cas-effector-protein complexes can function as monitors. Once foreign segments enter prokaryotic genomes, crRNA-Cas-effector-protein complexes can cleave them, inactivating the foreigners’ activity in the interference stage. The effectors of class 2 CRISPR-Cas systems employ a single protein, and such effectors can be conveniently manipulated; additionally, their guide RNAs are easily synthesized, which has led to the widespread use of class 2 systems in gene editing (7,8) and human essential gene screening (9,10). These systems are the best studied among all the recently discovered bacterial defense weapons (11). Moreover, the functions performed by CRISPR-Cas systems include not only anti-MGE immunity but also the modulation of other biological processes such as gene regulation, bacterial virulence regulation, and DNA repair (12,13).

The task of annotating and classifying Cas loci is important. For example, an available server can facilitate CRISPR-Cas searching and the identification of Cas-associated accessary proteins. Therefore, accurate assignment of the CRISPR-Cas (sub)type is necessary. To address the issue of annotating CRISPR, the scientific community has developed some user-friendly web servers and standalone software programs, such as CRISPRfinder (14), CRISPRDetect (15), PILER-CR (16), CRISPRdb (17), and CRISPRmap (18), which have perfectly solved CRISPR annotation and classification. For the annotation of Cas protein, Chai *et al.* constructed a web-based server, HMMCAS, to identify Cas genes (19). Users can submit only protein sequences for Cas protein detection in HMMCAS. However, HMMCAS cannot determine which Cas proteins constitute a Cas locus or what (sub)type a Cas locus belongs to. Zhang and Ye provided a web-based server, CRISPRone, that can perform Cas annotation (20). CRISPRdisco, an automated pipeline, was developed to discover and analyze CRISPR-Cas systems and can determine the type and subtype (21). CRISPRCasfinder, an updated version of CRISPRfinder, was developed (22) and can determine Cas proteins, Cas loci and (sub)type in addition to identifying CRISPR arrays. Recently, a knowledge base, CRISPRminer, was constructed to investigate the interaction between microbe and phage. In this work, the authors integrated useful annotation services such as CRISPR-Cas system classification module and anti-CRISPR proteins (Acrs) annotation module (23). Here, we have provided a freely available webserver, CasLocusAnno, to accurately and quickly determine Cas proteins, Cas loci and the corresponding (sub)type based on a previous work (1).

## MATERIALS AND METHODS

### General outline of methods and data downloading

We developed a web-based service program to perform Cas protein, Cas locus and (sub)type annotation basically following Makarova *el al*.’s method (1) with some adaptions. The procedure is schematically shown in Figure 1. To accelerate the annotation speed, we adopted multithreading in every annotation step to enable our server to finish the annotation as fast as possible. Multiple profiles can sometimes match different regions of the same protein, and it is important to choose an appropriate profile due to the correlation between the (sub)type assignment of a Cas locus and a Cas profile selection. To select an appropriate Cas profile when multiple profiles match different regions of the same protein, we used a graph search algorithm, the Markov cluster algorithm (MCL) (24), to solve the Cas profile selection problem. Thus, our server is good at identifying Cas proteins with multiple domains. We also proposed the concept of a <u>m</u>aximum <u>c</u>ontinuous <u>C</u>as <u>s</u>ubcluster (MCCS) to obtain a tightly clustered locus. When performing prediction, users can submit not only whole-genome nucleotide sequence but also whole-genome protein sequences. Users can also submit a RefSeq accession number for annotation. Thus, our server provides an expanded set of options for submitting data.

**Figure 1.**
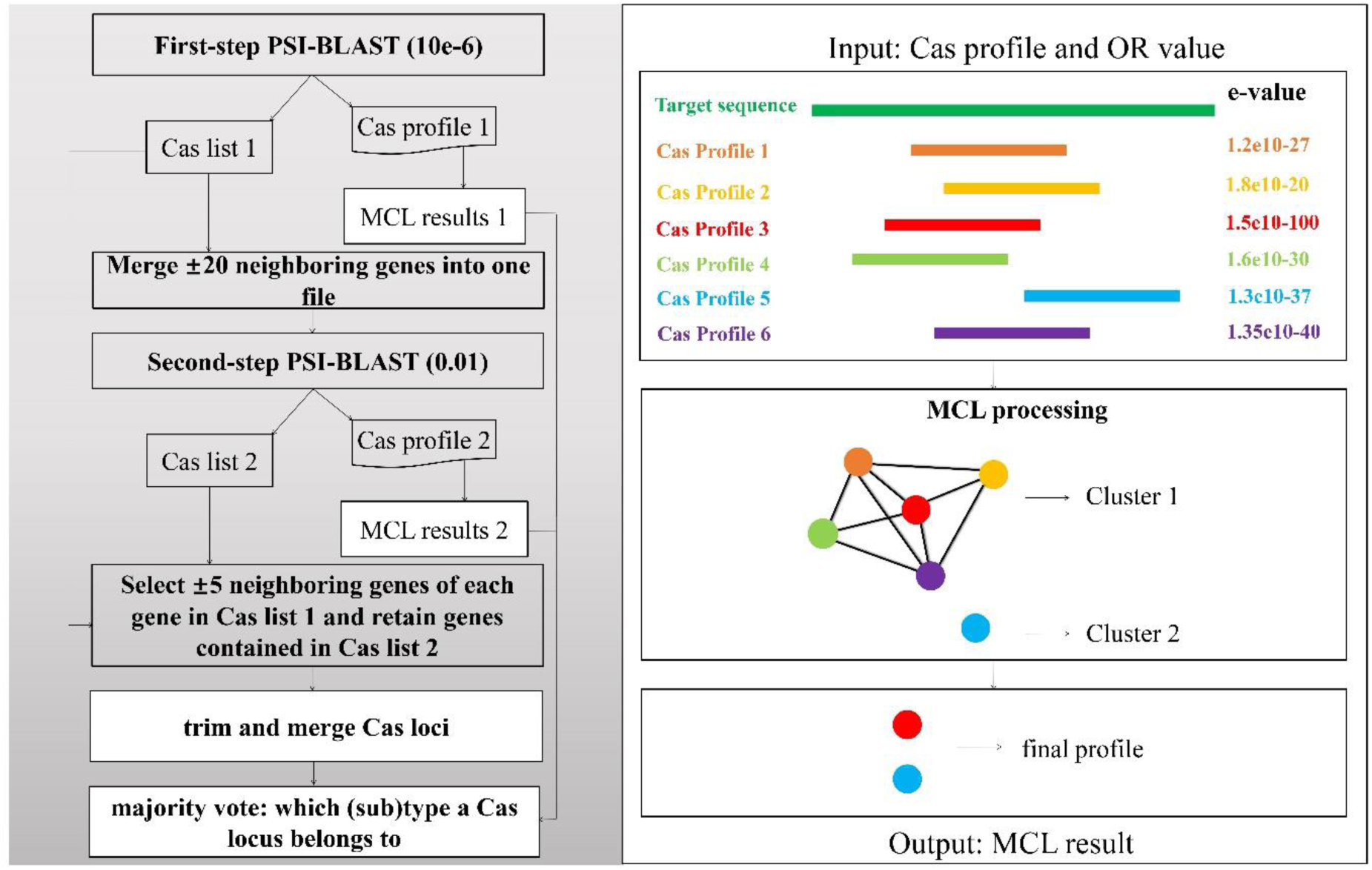
The workflow of annotating Cas loci and the principle of selecting an appropriate Cas profile. We perform the annotation procedure beginning with identifying Cas protein lists and followed by MCL clustering to handle the overlapping segments of the target position in a subject sequence. The Cas profiles obtained from the two PSI-BLAST steps are handled by MCL (right panel of the figure), and appropriate Cas profiles (MCL results) of targeting sequences are returned.

Cas profiles can be downloaded from ftp://ftp.ncbi.nih.gov/pub/wolf/_suppl/CRISPR2015/. The compressed package makarova_cas_NRM2015.profiles.tgz stores Cas profiles that were employed in Makarova *et al*.’s work (1). The compressed package makarova_cas_NRM2015-201604.profiles.tgz is an updated version of makarova_cas_NRM2015.profiles.tgz, which contains c2c1, c2c2, and c2c3 Cas profiles, therefore allowing the annotation of type VI subtypes.

### Identification of potential Cas proteins via two-step PSI-BLAST search

Several Cas genes cluster together to form a Cas operon or locus. A primary step in annotating the Cas operon is to identify Cas proteins. We downloaded Cas profiles (makarova_cas_NRM2015.profiles.tgz and makarova_cas_NRM2015-201604.profiles.tgz) from ftp://ftp.ncbi.nih.gov/pub/wolf/_suppl/CRISPR2015/, which were then used as queries to search Cas proteins in a subject bacterium. Similar to Makarova *et al.*’s method, a two-step PSI-BLAST (1) search was adopted. The first PSI-BLAST step was performed with an e-value of 10e-6 followed by selecting ±20 neighboring genes around each identified Cas protein. The PSI-BLAST search criteria are the same as in previous work (1), but the detailed procedure for handling these neighboring genes is different. We merged all the neighboring genes into a file. A single sequence may appear multiple times in the merged file if two Cas proteins are close to each other. Therefore, we termed sequences sharing with the same IDs as redundant sequences in this work and removed the redundant copies. All the nonredundant neighboring genes were included in the second PSI-BLAST search step with an e-value of 0.01. To accelerate annotation, we adopted multithreading in the PSI-BLAST searches. Thus, our server can finish the annotation in less than ∽28 seconds for chromosomes with ∽30 to ∽10500 protein sequences. Finally, two Cas protein lists could be obtained via the two-step PSI-BLAST search. Multiple profiles can sometimes match different regions of the same protein. It is important to choose an appropriate profile because of the correlation between the (sub)type assignment of a Cas locus and a Cas profile selection. In the next section, we used a graph search algorithm, MCL (24), to solve this issue.

### Selection of an appropriate Cas profile based on MCL in conflicting cases

To solve the issue of Cas profile selection, we defined a parameter *OR* (<u>o</u>verlapping <u>r</u>atio), which car be used to measure the degree of overlap between two optimal target segments within a subject protein sequence. The parameter *OR* is intrinsically distinct from the coverage ratio in the BLAST program although their calculated formulas appear to be the same. The *OR* reflects the overlap of two differen segments in the same subject sequence matched by different Cas profiles, whereas the coverage ratic reflects the alignment status between the query and subject sequences. The *OR* can be calculatec using the following equation:

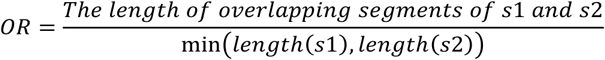

where *s1* and *s2* represent the segments matched by two different query Cas profiles in a subject sequence. It can be calculated from the length of the overlapping target segments matched by each profile pair divided by the smaller length of two target segments. Given that there are *N* Cas profiles matching one sequence, we can obtain *N(N-1)/2 OR* values. If the *OR* is higher than 90%, an edge will be formed between the two Cas profiles, and we assign the edge weight equal to 1, otherwise to 0. Using this method, we transform an alignment into a graph with edge weights of 1 and 0. MCL is employed to cluster the graph points according to the topology of the graph. The representative Cas profile with the minimal e-value within a cluster group is chosen as the final Cas profile. Two Cas protein lists and their corresponding Cas profiles are obtained via two-step PSI-BLAST searches and MCL, respectively. We next determine the Cas locus and its (sub)type by expanding an MCCS region and a voting strategy.

### Determination of Cas locus, (sub)type via expanding an MCCS region and voting procedure

Makarova *et al.*’s strategy was adopted to determine a Cas locus (1), but we also added some adaptions. Because some false positive Cas proteins may be found in the second-step search, we may provide an incorrect Cas operon by selecting ±5 neighbor genes identified in both steps of the PSI-BLAST searches. Thus, we merely select ±5 genes around each Cas gene obtained in the first of the PSI-BLAST search and rule out sequences that are not contained in the Cas gene list obtained from the second PSI-BLAST search. Any gene belonging to the list will be retained. Finally, potential Cas loci that share the same sequence IDs are trimmed and merged according to the previous work (1). Cas proteins in a potential Cas locus are clustered tightly to perform their functions. MacSyFinder depends on colocalization rules to model Cas clusters (25). Here, we introduced another concept, MCCS, to further refine the potential Cas locus. MCCS refers to a subcluster that has the maximum number of consecutive Cas proteins, which means that there are no gaps between each Cas protein pair, and the Cas protein number in an MCCS should be more than 3. Figure 3 shows an example of how a MCCS region is expanded. First, a potential Cas cluster is classified into several subclusters according to the continuity of Cas proteins. Second, MCCS is chosen according to our definition. Finally, an MCCS is expanded according to the distance between the MCCS and the subcluster located upstream or downstream, as well as the Cas protein number of the subcluster. If the distance is equal to 2, which means that there is only one gap protein between each boundary Cas protein pair in the two subclusters, and the neighboring subcluster of the MCCS has more than two Cas proteins, we expand the MCCS region until it does not meet the expanded criteria that we set. The originally predicted Cas proteins that are not added into a MCCS region during its expansion will be discarded.

**Figure 3.**
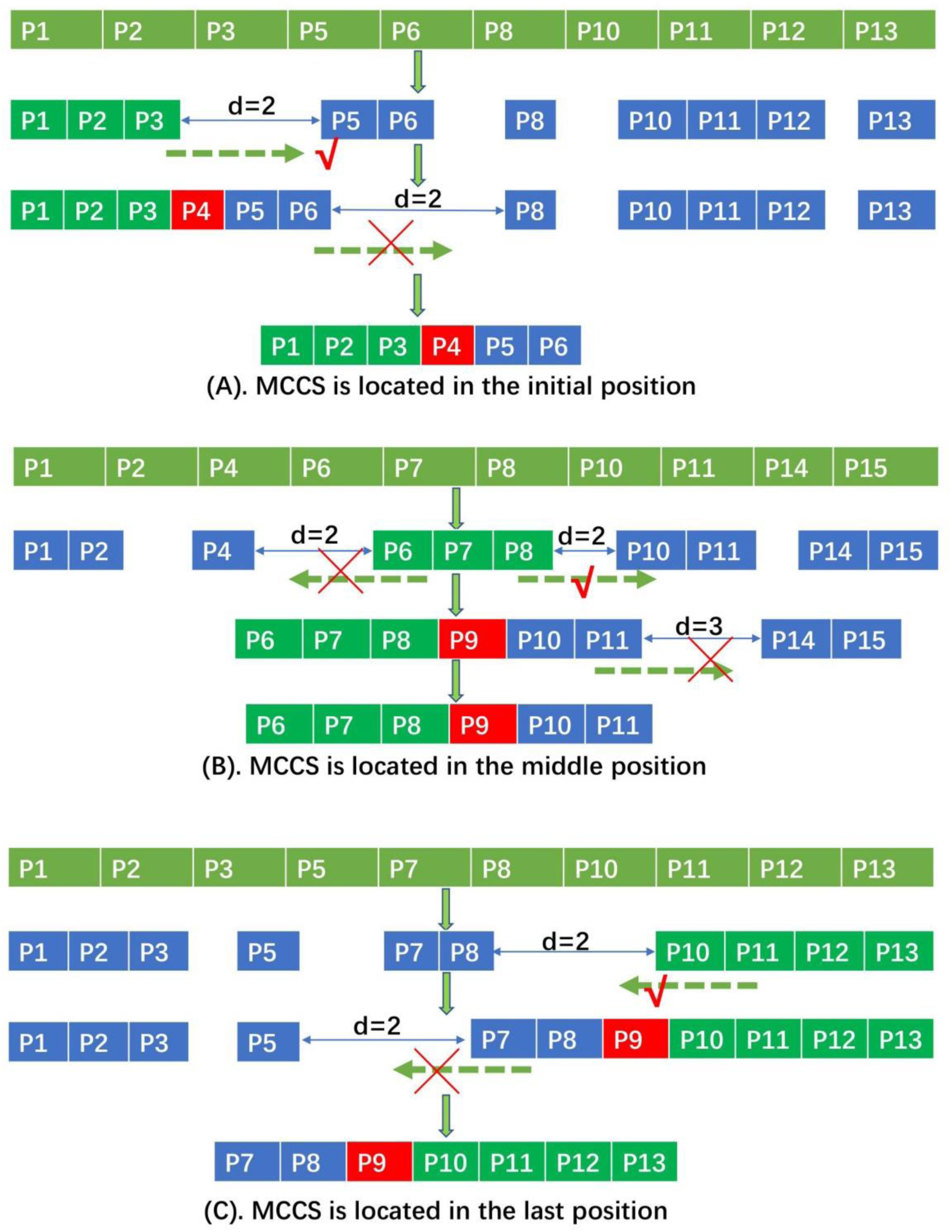
Examples to illustrate how an MCCS is expanded. The P*_index_* represents Cas protein, and the successive index on the right side of *P* denotes neighboring Cas proteins. Consecutive proteins with a light green background correspond to the MCCS region. A square with a red background represents a potential Cas protein added during the expansion procedure. The symbol “√” denotes a MCCS that satisfies the expansion conditions. The symbol “×” denotes an MCCS that does not satisfy the expansion conditions. (A) indicates that the MCCS is located at the initial position, and we will expand it from the initial to the last position; (B) indicates that the MCCS is located at the middle position, and we will expand it from the middle to the initial and last position, respectively; and (C) indicates that the MCCS is located in the last position, and we will expand it from the last to the initial position.

We performed the voting procedure beginning with a final Cas locus obtained from the MCCS expanding procedure. A potential Cas locus is labeled “not a Cas locus” if the Cas protein number is less than 3, and its information is not be displayed on our server page; otherwise, a potential Cas locus will enter the next judgment process. A potential Cas locus is divided into “not a valid Cas locus” and “valid Cas locus” according to whether it contains effector Cas proteins or not. The latter case will enter the final judgment process. A potential Cas locus can be determined by its signature effector or voting strategy. If a subtype and its corresponding type appear in the same locus, the type will be regarded as the subtype during the voting decision process.

## RESULTS

### Accuracy and speed assessment

To systematically evaluate the speed and accuracy of CasLocusAnno, we downloaded all whole protein sequences located on 1263 chromosomes and plasmids that were used by Makarova *et al.* in their work (1) from NCBI (<u>n</u>ational <u>c</u>enter for <u>b</u>iotechnology <u>i</u>nformation). The protein sequence number in the selected chromosomes ranges from ∽30 to ∽10500. We used CasLocusAnno to determine Cas proteins, Cas loci and their corresponding (sub)types in each chromosome. Figure 2(a) displays the distribution of each type and (sub)type in our annotations. Obviously, type I systems are more widespread in bacterial and archaeal genomes than others. As we can see, ∽66% of genomes have type I systems, which is basically consistent with Makarova *et al.*’s results (64% in bacterial genome, 60% in archaeal genomes). Type IV and type V systems represent only ∽2.03% (∽1.627% for IV system, ∽0.4068% for V system) among all types. This result is also consistent with Makarova *et al.*’s findings (∽2% overall). We excluded some loci with two Cas proteins; therefore, CAS-I-B is the most abundant subtype system, followed by subtypes I-E and I-C. This conclusion differs slightly from that obtained by the previous method, in which the most abundant CRISPR-Cas system is subtype I-E, followed by I-B and I-C. For type II systems, CAS-II-C is the most abundant, followed by subtype II-A and II-B systems. Figure 2(b) displays the running time under different protein numbers on each chromosome. We adopted multithreading to perform Cas protein, Cas locus and (sub)type annotation; thus, our server can complete a prediction within ∽28 seconds for the whole protein sequence submission of a chromosome. Considering that CRISPRone needs more than 15 mins to finish Cas protein annotation for a 5 Mb genome, CRISPRCasFinder needs 1∽3 mins, and CRISPRDetect needs 2∽20 mins (refer to supplementary material 2 in reference 22), we consider 28 seconds to be reasonably acceptable.

**Figure 2.**
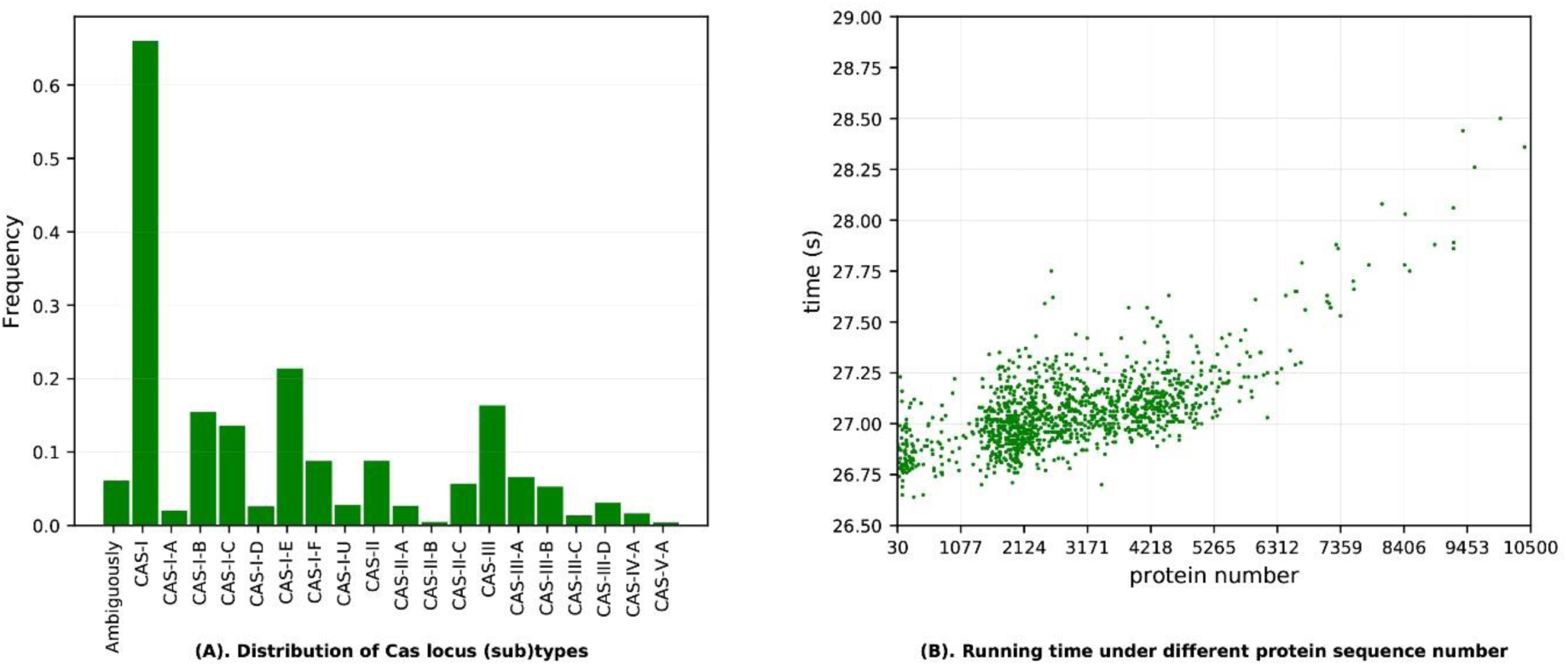
(A) Distribution of each (sub)type among our annotations. (B) Running time for chromosomes with different protein numbers. The results indicate that our server can finish annotation ∽28 seconds.

Sometimes multiple profiles can match different regions of the same subject protein (refer to left panel in Figure 1). It is important to choose an appropriate profile due to the correlation between the (sub)type assignment of a Cas protein and a Cas profile selection. In our work, we used MCL to solve this issue. A graph reflecting the *OR* between different regions matched by each pair of Cas profiles is constructed and clustered by MCL according to edge weight and graph topology. In each group, the profile with the lowest e-value is chosen. Thus, our selected profiles are reasonable. For example, the I-F subtype in NC_005966 has a Cas6f protein, which matches profile pfam09618 with an e-value of 1. 37e-10 in our work; however, the same protein corresponds to cd09739 with an e-value of 2.08e-10 in Makarova *et al.*’s work. Another example is the I-E subtype in chromosome NC_009484, where gi|148260808|ref|YP_001234935.1| matches cd09639 with an e-value of 3.29e-53 in Makarova *et al.*’s work, whereas its profile is C0G1203 in ours with an e-value of 2.76e-53, which is a lower e-value than the former. CasLocusAnno can always select the profile with the lowest e-value in each MCL group; therefore, it can select a more suitable Cas profile.

### Cas protein comparison with CRISPRCasFinder and MacSyFinder

Users need to submit whole-genome nucleotide sequences when using CRISPRCasFinder to perform Cas annotation. CRISPRCasFinder utilizes Prodigal (version 2.6.3) to identify protein sequences (26), and CRISPRCasFinder then utilizes MacSyFinder (version 1.0.5) (25) combined with updated Cas profiles to determine Cas proteins, Cas loci and their corresponding (sub)types. For comparison, we selected 943 chromosomes that are well annotated, where each locus in those chromosomes contains more than two Cas proteins in Makarova *et al.*’s supplementary material. Considering that the whole protein sequences are easily obtained for the selected chromosomes, it is not necessary to re-annotate their coding sequences by Prodigal. We used MacSyFinder (version 1.0.5) and the updated Cas profiles, which were obtained from a standalone version of CRISPRCasFinder, to identify Cas proteins and Cas loci and (sub)types. Figure 4 illustrates the overlapping and nonoverlapping Cas protein numbers annotated by CasLocusAnno, Makarova *et al.*, and MacSyFinder. CasLocusAnno, Makarova *et al.*, and MacSyFinder identified a total of 8447, 9327 and 7939 Cas proteins in all Cas loci, respectively. We used Makarova *et al.*’s annotation results as a benchmark and calculated the accuracy and <u>a</u>dditional <u>p</u>rediction <u>r</u>ate (APR) using the following equations:

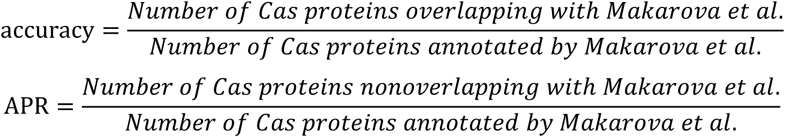

**Figure 4.**
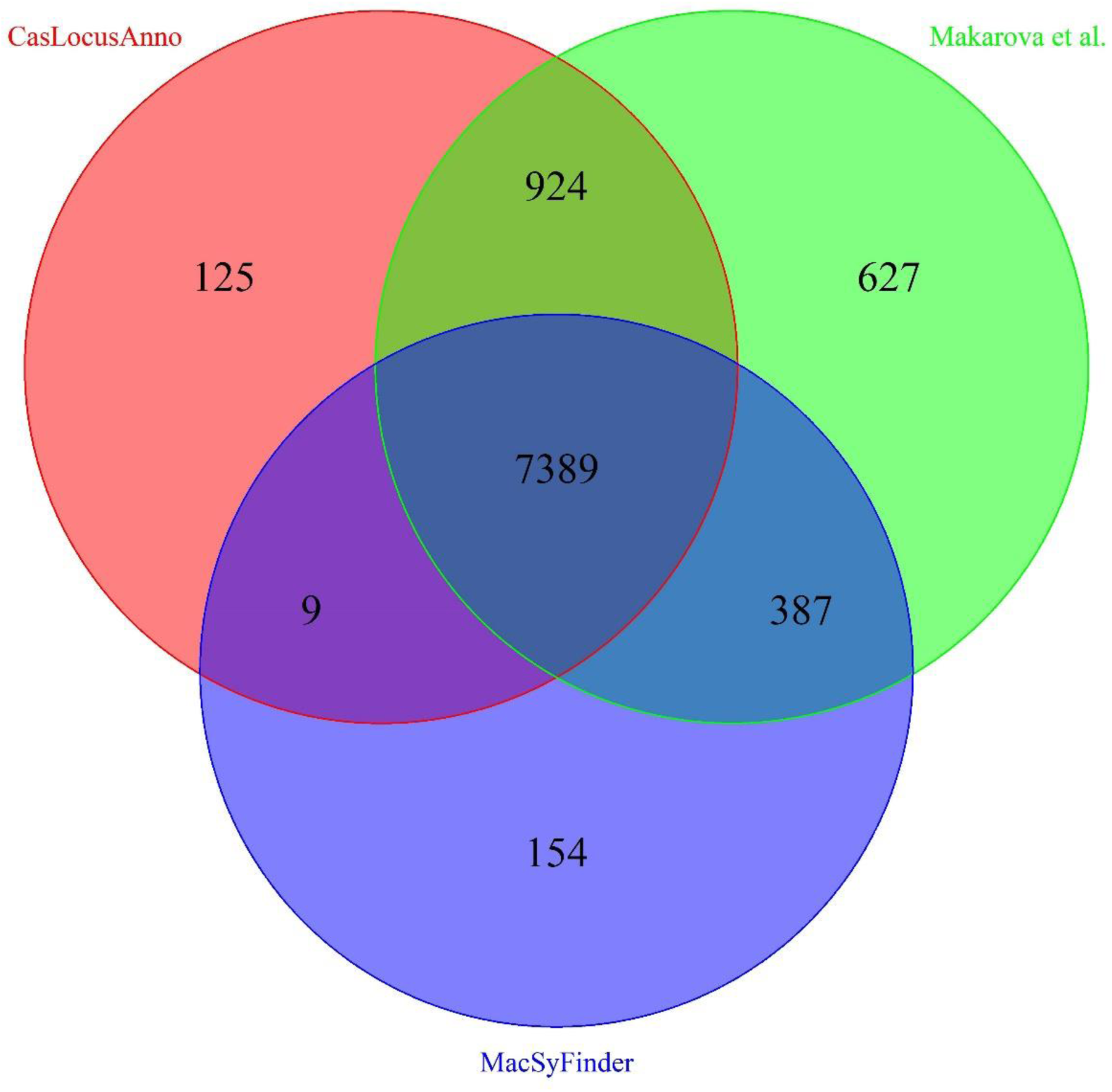
compares Cas proteins annotated by CasLocusAnno, Makarova *et al.*, and MacSyFinder for 943 selected chromosomes. The results annotated by Makarova *et al.*, CasLocusAnno, MacSyFinder are represented by circles with red, green and blue backgrounds, respectively.

CasLocusAnno and Makarova *et al.*’s results share a total of 8313 Cas proteins over all Cas loci, whereas MacSyFinder and Makarova *et al.*’s results share only 7776 Cas proteins. Our accuracy is ∽89.13% (8313/9327), whereas the accuracy of MacSyFinder is ∽83.3% (7776/9327). In addition, CasLocusAnno identified 134 additional Cas proteins, and MacSyFinder identified 163 additional Cas proteins. Thus, our APR is ∽1.4% (134/9327), whereas MacSyFinder’s additional prediction rate is ∽1.7% (163/9327). Therefore, the performance of CasLocusAnno is significantly better than that of CRISPRCasFinder and MacSyFinder. The annotation results of CasLocusAnno and MacSyFinder can be downloaded from the download page of the CasLocusAnno server. MacSyFinder is an excellent software program for mining genomes for molecular systems (25). It selects the profile with the highest score when faced with situation in which a query protein is matched by multiple profiles. If a protein has multiple domains, such as a Cas protein fused by other Cas proteins, it is difficult for MacSyFinder to detect because it selects only the profile with the highest score and ignores others. CasLocusAnno employs MCL to select a profile, which can avoid such cases to some degree.

### CasLocusAnno allows users to submit data via three methods

To facilitate researchers’ work, we have allowed CasLocusAnno to accept multiple types of data. Users can submit the whole protein sequences of a chromosome or the whole nucleotide sequence of a chromosome. For genome nucleotide sequence submission, the primary step is to determine all protein-coding genes via ZCURVE 3.0 (27), then detect the Cas locus and (sub)type. For some well-annotated genomes, users can submit only the whole protein sequences of a chromosome. The RefSeq accession number is also acceptable.

## CONCLUSION

CasLocusAnno can be used as an alternative supplementary web-based software to existing methods such as CRISPRCasFinder, MacSyFinder, HMMCAS, and CRISPRdisco. CasLocusAnno offers some advantages. It can select a more reasonable Cas profile because in every cluster, it can always choose a profile with the minimum e-value, which leads it to accurately confer a correct (sub)type. CasLocusAnno can identify fused Cas proteins; however, MacSyFinder cannot detect such proteins because it selects only the profile that has the highest score. Our server can accept whole protein sequences, whole nucleotide sequence or the Ref accession number of a chromosome to perform annotation. Compared with two existing excellent software packages, CRISPRCasFinder and MacSyFinder, CasLocusAnno can identify Cas proteins with higher accuracy and lower APR. Finally, CasLocusAnno can return annotation results in less time due to the advantage of parallel computing.

## DISCUSSION

Since CRISPR-Cas technology has brought unprecedented changes to gene editing and gene regulation, it is crucial to improve the efficiency and specificity of existing CRISPR-Cas systems and discover more efficient and special new Cas proteins. Some specific Cas proteins have been discovered, including Cas9 targeting dsDNA, Cpf1 targeting dsDNA, C2c2 targeting ssRNA (2) and Cas14 targeting ssDNA (28). Smargon *et al.* developed a computational pipeline to determine putative CRISPR-Cas loci without the most conserved Cas1 and Cas2 proteins among bacterial genomes, which helped validate Cas13b as a type VI-B system that cleaves target RNA and displays RNase activity (29). Thus, an available server software can help researchers identify other Cas effector proteins. Some ancillary Cas proteins exist in addition to the core proteins in Cas arrays. Identifying Cas loci and (sub)types can facilitate the search for Cas-associated accessory proteins. For example, Shmakov *et al.* systematically determined Cas-associated genes by determining CRISPR-Cas systems and (sub)types (30). Recently, Shah *et al.* identified some co-functional accessory genes within the type III CRISPR-Cas system using a guilt-by-association approach based on Makarova *et al.*’s Cas loci data (1).

Although, some useful web-based servers (22,23) and standalone implemented programs (21) have been emerging and can be available from network and GitHub (31) for CRISPR-Cas filed, our constructed service can be still as a complementary resource mainly because of the following reasons:CasLocusAnno can select a more reasonable Cas profile because in every cluster, it can always choose a profile with the minimum e-value, which leads it to accurately confer a correct (sub)type. CasLocusAnno can identify fused Cas proteins; however, MacSyFinder and CRISPRCasFinder cannot detect such proteins because it selects only the profile that has the highest score. Our server can accept whole protein sequences, whole nucleotide sequence or the Ref accession number of a chromosome to perform annotation.

<u>S</u>elf-<u>t</u>argeting (ST) means that a spacer and its targeting segment appears in the same prokaryotic organism. ST can lead to reduced bacterial fitness. Rauch *et al.* scanned 275 *Listeria monocytogenes* genomes to discover which strains exhibit ST and used ST as a biological marker to screen for bacteria bearing Acr (32). Recently, researchers utilized ST to discovery several Acrs that can inhibit the activity of Cpf1 protein (31). Almost at the same time, Marino *et al.* also descripted several Acrs that can inhibit the activity of Cpf1 protein (33). An Acr can also be classified according to the CRISPR-Cas (sub)type that it inhibits. Furthermore, Cas locus and (sub)type annotation can enrich the annotation of functional elements in bacterial genomes in addition to annotating protein-coding genes, genome islands, essential genes, etc. The diverse CRISPR-Cas systems imply that phages should bear multiple Acrs to inhibit activity of the host immune system. Inspired by Rauch *et al.*’s perspective, we speculate that if a bacterium satisfies two conditions, the presence of ST and an essential targeting gene, then the bacterium would have a higher likelihood of containing Acrs. For the essential gene prediction issue, we have developed Geptop (34), and for Cas locus annotation, we have developed CasLocusAnno. We will continue our work and increase the overall number of Acr entries in Anti-CRISPRdb (35) and the unified resource of Acrs in a Google document (https://tinyurl.com/anti-CRISPR) (36).

## SUPPLEMENTARY DATA

Supplementary Data are available at //cefg.uestc.edu.cn/CasLocusAnno/download.html.

## ACKNOWLEDGMENTS

The authors thank Hong-Li Hua for kindly making suggestions about our server. Abraham A. Labena kindly made English language revision. Dr. Yuri I. Wolf kindly answered our questions about their annotation pipeline via email.

